# Prevalence of Diversified Antibiotic Resistant Bacteria within Sanitation Related Facilities of Human Populated Workplaces in Abbottabad

**DOI:** 10.1101/2020.05.05.078485

**Authors:** Jawad Ali, Malik Owais Ullah Awan, Gulcin Akca, Iftikhar Zeb, Bilal AZ Amin, Rafiq Ahmed, Muhammad Maroof Shah, Rashid Nazir

**Affiliations:** Department of Biotechnology, COMSATS University Islamabad (CUI), Tobe Camp, University Road, postal code 22060, Abbottabad Campus, KPK Pakistan; Department of Environmental Sciences, COMSATS University Islamabad (CUI), Tobe Camp, University Road, postal code 22060, Abbottabad Campus, KPK Pakistan; Department of Medical Microbiology, Faculty of Dentistry, Gazi University Ankara, 8.cad.82.sok.No:4 06510 Emek Çankaya Turkey

**Author notes:** Correspondence: Rashid Nazir, PhD.

**Keywords:** Hygiene, sanitation, antibiotics, antibiotic resistance, workplaces, antibiotic resistant bacteria

## Abstract

Antibiotics discovery was a significant breakthrough in the field of therapeutic medicines, but the over (mis)use of such antibiotics (n parallel) caused the increasing number of resistant bacterial species at an ever-higher rate. This study was thus devised to assess the multi-drug resistant bacteria present in sanitation-related facilities in human workplaces. In this regard, samples were collected from different gender, location, and source-based facilities, and subsequent antibiotic sensitivity testing was performed on isolated bacterial strains. Four classes of the most commonly used antibiotics i.e., β-lactam, Aminoglycosides, Macrolides, and Sulphonamides, were evaluated against the isolated bacteria.

The antibiotic resistance profile of different (70) bacterial strains showed that the antibiotic resistance-based clusters also followed the grouping based on their isolation sources, mainly the gender. Twenty-three bacterial strains were further selected for their 16s rRNA gene based molecular identification and for phylogenetic analysis to evaluate the taxonomic evolution of antibiotic resistant bacteria. Moreover, the bacterial resistance to Sulphonamides and beta lactam was observed to be the most and to Aminoglycosides and macrolides as the least. Plasmid curing was also performed for MDR bacterial strains, which significantly abolished the resistance potential of bacterial strains for different antibiotics. These curing results suggested that the antibiotic resistance determinants in these purified bacterial strains are present on respective plasmids. Altogether, the data suggested that the human workplaces are the hotspot for the prevalence of MDR bacteria and thus may serve the source of horizontal gene transfer and further transmission to other environments.

## Introduction

An antibiotic is a substance produced (synthetically or mostly) by an organism to kill or inhibit the growth of another organism [1]. Antibiotics can mainly be classified based on their molecular structure [2] and mode of action [3] and so, they are beta lactam, macrolides, Aminoglycosides and Sulphonamides. Moreover, the very first antibiotic discovered, i.e. penicillin is a beta lactam [4].

Though, antibiotic discovery was one of the most significant breakthroughs in the field of therapeutic medicines enabling the treatments of serious bacterial infections [5] and thus crucially reducing the morbidity and mortality across the globe [6]. However, soon after the introduction of antibiotics in the clinical settings, the microbes also developed different strategies and mechanisms to overcome the effects of these antibiotics, cumulatively called antibiotic resistance (AR) [7]. Such resistance may develop due to the mis/overuse of antibiotics in treating the infections and also because of the antibiotics’ frequent use in agriculture, aquaculture and veterinary practices [8]. Along with antibiotics resistance development, these factors may also contribute significantly to resistance spread in the environment [9]. The overuse of antibiotics, poses a selection pressure under which the susceptible bacterial strains are eliminated while resistant ones survive and even transfer their resistant capability further to other bacterial (susceptible) strains [10]. Consequently, the number of antibiotic resistant bacteria are increasing over time. Ultimately, the antibiotic resistant bacterial (pathogenic) strains can cause serious health issues to human and animals [11].

Bacteria have adopted various molecular strategies to become resistant to antimicrobial drugs [12; 13] and, thus, bacterial antibiotic resistance has been developed via two main mechanisms i.e., genetic mutations and resistant gene (transfer) acquisition [14]. These antibiotic resistance determinants can prevail in microbial communities and even be transferred in the environment from one point to another, and such transfer from one bacterium to another is normally termed as horizontal gene transfer (HGT). Mobile DNA can move from one part of the genome to another or between different genomes, and antibiotic resistant genes are generally present on such mobile DNA e.g., plasmids and transposons [15]. Noticeably, the plasmids are extrachromosomal DNA, replicating independently from the host chromosome and are an essential source of HGT. While, conjugation, transformation, and transduction are the main mechanisms of horizontal gene transfer in bacteria.

Along with biological and molecular factors, the environment is also playing a significant role in the spread of antibiotic resistance. One of the environmental factors which contribute to multi-drug resistance is sanitation and non-hygienic conditions. Specifically, the improper use of antibiotics causes their residues and metabolites to prevail in human and animal wastes i.e., in water, soil, and water-dependent food crops and consequently the bacteria present in these environments (and exposed to these antibiotics’ effect) may acquire antibiotic resistance. For instance, the wastewater treatment plants are known as hotspots for antibiotic resistance and spread [45]. Water from various sources flows into these treatment plants where a variety of bacteria and resistance genes are present, which may also transfer from one bacterium to another and further contribute in the environmental spread of drug resistance [1]. The sanitation and hygienic conditions in Pakistan are deplorable, which may significantly contribute to the spread of pathogenic bacteria and infectious diseases to the people living (and/or working) in such environments. Such unhealthy hygienic practices contribute to the spread of antibiotic resistance and, ultimately, the resistant strains of bacteria [16].

Moreover, humans can also contribute in drug resistance, as the intestine contains a diversity of bacteria, which can potentially be a source of antimicrobial resistance. For example, extra intestinal pathogenic *E. coli* is measured as most critical contributor to antibiotic resistance [17]. Furthermore, different people are exposed to different environments and different microbiota (e.g., pathogens) and when they interact in a common workplace, there is a higher chance of bacterial and ARGs’ transfer to each other [18]. In these environments, different commensal bacteria are also present which themselves are non-pathogenic but can develop resistance via interacting with other antibiotic resistant bacteria and may even transfer the resistance genes to pathogenic bacteria [16]. In this regard, the Abbottabad region of KPK province in Pakistan is a densely populated area with inappropriate hygienic management and so presents a suitable situation to monitor the existence of drug resistance within human populated workplaces. In this study, we, therefore, have evaluated different educational institutes of Abbottabad to assess the prevalence and spread of antibiotic resistance in the context of existing hygienic conditions. Educational institutes have a high human flow, and so the chances of AR spread are more because a large number of (commensal as well as pathogenic) bacteria are present in such environments. The transfer potential of AR genes in these situations is greatly enhanced, subsequently leading to the production of multi-drug resistant pathogenic bacteria. The overall purpose of this study was to isolate and identify the bacteria showing resistance to multiple antibiotics and also to characterize them for antibiotics susceptibility. The specific objectives were, (1) Isolation, identification, and characterization of multi-drug resistant bacterial strains from populated human workplaces of Abbottabad, and (2) Plasmid curing of bacterial strains, to evaluate the drug resistance mechanisms, isolated from various sanitation-related environmental samples.

## Materials and Methods

### Sample Collection

Samples from sanitation facilities were collected from three different human-populated locations in Abbottabad Pakistan i.e., COMSATS University campus (A-block), Ayub Medical College (AMC), and Ayub Teaching Hospital (ATH). The sampling layout is illustrated in figure 1. Specifically, the samples were collected from places with high chances of bacterial presence i.e., washrooms. Sludge samples were collected (in sterilized glass vials containing autoclaved water) from the washrooms’ basins and sanitary pots via using sterilized swab sticks. Moreover, the male and female washrooms were sampled separately. Three (turbid aquaculture) (sub)samples from each location and object were sampled to make a composite representative sample, and this way, the three independent biological replicates were collected for each treatment scenario. The samples were then transported to the laboratory in COMSATS in a thermos-flask packed with ice to keep the biological/chemical material of the samples intact and, to perform the subsequent necessary analyses.

**Figure 1.**
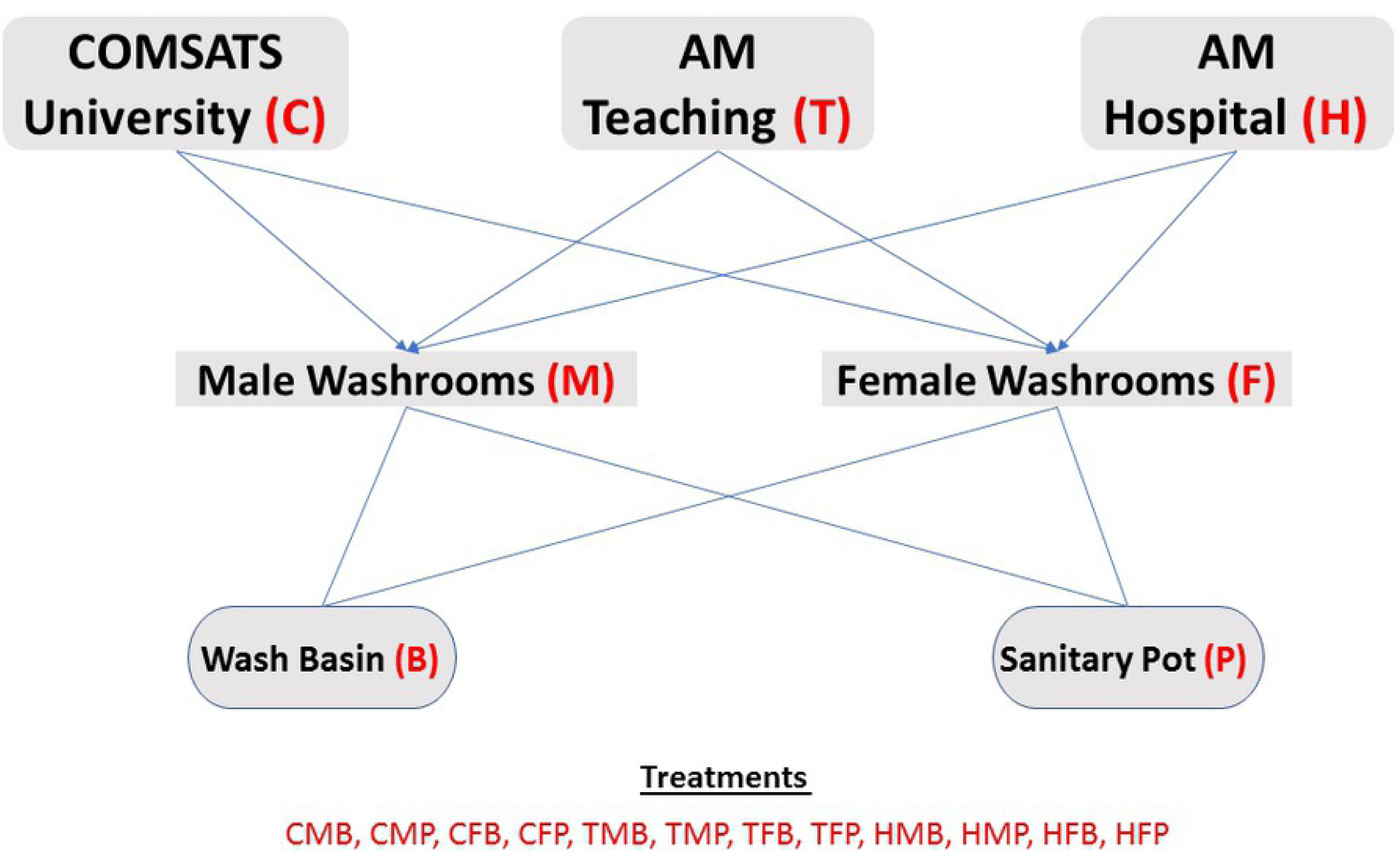
Schematic description of the sampling plan executed in this work, to evaluate the antibiotic resistant bacterial load present in hygiene related scenarios of populated human workplaces of Abbottabad, PK.

### Purification of hygiene-related Bacterial Isolates

The cultivable bacteria were cultured on Luria-Bertani agar (LBA, Sigma-Aldrich, Germany) plates. For this purpose, serial dilutions of the hygiene-related sludge samples were performed, and 100µl of the appropriate dilution was spread on LBA plates, consequent to the incubation at 37° C for 24-48 hours [19]. After the incubation, plates were observed for bacterial growth and colony forming units (CFU) were counted for each plate. After evaluating the initial bacterial growth, different colonies were selected (based on their distinct morphology) for purification through the streak plat method [20].

Fresh LBA plates were prepared, and each selected bacterial strain was inoculated for proper growth until the appearance of purified monotonous colony type. The single purified colonies were then used for further experiments. Moreover, all the purified bacterial strains were grown in Luria-Bertani broth (LB, Sigma-Aldrich, Germany) and preserved in glycerol solution to store as −80°C stock.

### Antibiotic susceptibility testing for purified bacterial strains

For hygiene-related isolated bacterial strain, antibiotic susceptibility tests were performed for eight commonly used antibiotics of <u>four different chemical classes</u> i.e., β-Lactams, aminoglycosides, macrolides, and Sulphonamides (Table 1). For this purpose, fresh LBA plates were prepared via mixing the sterilized aliquots of test antibiotics in the autoclaved medium [21] at a concentration of 100µg/ml. Moreover, the bacteria were also inoculated on LBA plates with no antibiotic and were used as a control for microbial growth under the same incubation conditions.

**Table 1:** Antibiotics used in this study along with their chemical classification and established mode of action.

| Name of Antibiotics | Antibiotic class | Mode of action |
| --- | --- | --- |
| Co-amoxiclav<br>Ampicillin<br>Amoxicillin | β-Lactams | Inhibition of bacterial cell wall synthesis |
| Amikacin<br>Gentamicin | Aminoglycosides | Inhibition of bacterial cell membrane / protein synthesis |
| Clarithromycin<br>Azithromycin | Macrolides | Inhibition of bacterial protein synthesis |
| Co-trimoxazole | Sulphonamides | Inhibition of bacterial folate synthesis |

The isolated pure bacterial strains were inoculated on each of the antibiotic assisted LBA plates and incubated at 30°C. After 24 hours, the plates were observed for the presence of bacterial growth. According to this antibiotic susceptibility test, performed on agar plates, the inoculated bacterial strains growing on an antibiotic assisted plate were considered as resistant to that particular antibiotic and were also selected for further evaluations. While the bacteria not growing (even after 3-days) on antibiotic assisted LBA, though growing on non-antibiotic LBA (control) plates, were recorded as the susceptible one for that particular antibiotic.

### Molecular analyses

#### DNA Extraction

DNA was isolated from multi-drug resistant culturable bacterial strains as well as from culture-independent initial environmental (sludge) samples. For bacterial strains, for instance, the cultures were grown overnight in nutrient broth at 37°C, subsequently centrifuged at 13,000 rpm for 1 minute. The supernatant was discarded, and resultant cell pellet was re-suspended in 600 µl of lysis buffer [SDS (sodium dodecyl sulfate), proteinase K and Tris-EDTA (TE)]. Then, it was shaken to homogenize by vortex mixing and incubated for 1 hour at 37°C. Afterwards, phenol: chloroform was added and mixed until the homogeneous mixture was formed. The centrifugation was done at 13,000 rpm for 5 minutes to establish two separable layers. The upper layer containing DNA was transferred into another tube. An equal amount of 3M sodium acetate was added, mixed well and centrifuged at maximum speed for 5 minutes. The aqueous (transparent) layer was transferred to another tube. Chilled isopropanol was then added to the mixture and gently mixed. The tube was kept at −20°C for one hour, centrifuged at 13,000 rpm for 10 minutes, and the supernatant was removed. DNA pellet was then washed with 1ml 70% ethanol [centrifugation at 13,000 rpm for 2 minutes), and precipitated DNA was air-dried, and resuspended in TE buffer [22]. Each sample’s DNA was evaluated on agarose gel for purity and concentration.

#### Polymerase Chain Reaction

To identify the unknown bacterial strains, the 16s rRNA gene was amplified through PCR. The universal primers 27F (5’AGA CTC TCC TGA TGG GTT AG 3’) and 1492R (5’ACG TTA TTG CGA ACC GCT CTT 3’) were used for gene amplification [22] . The PCR ingredients for one reaction contained 1.5µl of extracted DNA, 12.5µl of the 2X ThermoScientific PCR master mix, 1µl of forward and reverse primers each and nuclease free water to make the volume up to 25µl.

After the reaction preparations, the PCR was conducted in a PTC-100 thermocycler as follows: denaturation at 94°C for 3 minutes and then 30 cycles at 94°C for 1 minute (denaturation), 56°C for 1 minute (annealing) and 72°C for 1 minute (elongation). The amplification products were analyzed on 1% agarose gel electrophoresis in TBE buffer (Tris-base, boric acid and 2mM EDTA) [14].

### DNA Sequencing, Basic Local Alignment Search Tool (BLAST) and Phylogenetic Analyses

The DNA amplified through PCR was purified and used for DNA sequencing. The ingredients used in sequencing reaction were DNA polymerase, primer (27F), four nucleotides (dATP, dTTP, dCTP, dGTP), fluorescent tags and DNA template. After the acquisition of DNA sequences, data was manually validated with Chromas (version 2.6.5). The resulting sequences were compared with those already present in the database, and bacteria were thus molecularly identified [23]. This BLAST analyses of bacterial 16s rRNA gene sequences were performed via online NCBI tools.

Based on the coverage and percent identity to our sequences, the similar bacterial sequences were downloaded from the NCBI database and used as reference sequences for subsequent phylogenetic analysis for our hygiene-related bacterial isolates. After the multiple sequence alignment, with MEGA (version X), a phylogenetic tree was constructed, which showed the evolutionary relationship between the isolated bacterial strains and the reference sequences obtained from the genomic database. The newly acquired hygiene-related bacterial sequences were deposited with accession numbers xxx – yyy in open access genomic database for public future use.

### Plasmid curing for hygiene-related bacterial strains

The plasmid curing was performed by the protocol developed earlier [24]. Briefly, the LB medium was prepared, sterilized by autoclavation, and acridine orange (filter sterilized) was added at a concentration of 0.1mg/ml. Then 5ml of this medium was poured into the test tube, and antibiotic resistant bacteria were inoculated into separate test tubes. It was then incubated at 30°C and 175 rpm for 24 hours. After 24 hours, the growth cultures were ten times diluted in fresh medium. Similarly, after 1, 2, 3, 5, and 7 days of incubation, the grown bacterial cultures were serially diluted and spread onto (respective) antibiotic assisted LBA plates. The strains not growing on these antibiotics assisted LBA, but growing on normal LBA, were considered as cured (plasmid losing) bacterial strains. The cured bacterial strains were also tested for antibiotic susceptibility profile (as described above). The colonies still growing on antibiotic assisted LBA (resistant ones) indicated that they did not lose plasmid via acridine orange and/or their resistance was not plasmid-borne (but is present on chromosomal DNA).

### Statistical Analyses

All the statistical analyses were performed in at least triplicates, and the obtained results are presented as averages. The data variations are expressed as standard deviations, shown in text (mostly in brackets) as numeric values and in graphs as error bars. For the assessment of cultivable bacteria, the CFU data were transformed into Log units per ml of the sampled sludge, and averages were presented in the graphs. ANOVA, t-test, and Tukey’s tests were performed (where applicable) to obtain the level of significance in variations of data. For the bacterial antibiotic resistance profiling, “R” studio analysis software was used. A dendrogram was constructed through cluster analysis in which the bacteria were grouped according to their antibiotic resistance to common antibiotics. DNA sequence analysis softwares (i.e., Chromas (v2.6.5) and MEGA X) were used for the validation and subsequent alignment analyses for the bacterial sequences.

## Results

### Cultivable Bacterial Abundance and diversity within Sanitation Facilities of Human Populated Workplaces

After serial dilution, sanitary samples were spread on LBA plates and incubated for 24 hours. Bacterial growth was observed on each plate, and colony forming units (CFUs) were enumerated. Figure 2a shows the Log CFU/ml of the samples collected from COMSATS University, Ayub Medical College, and Ayub Teaching Hospital. The CFU value of majority samples was in the range of 10^7^ - 10^8^ CFU/ml except for the two samples i.e., CFB (COMSATS Female washroom sanitary basin) and CFP (COMSATS Female washroom sanitary pot) where the CFU values were significantly lower than the others i.e., 10^6^ and 5×10^6^ CFU/ml for CFB and CFP respectively while there was no significant difference for total cultivable bacterial abundance for other two sampled locations neither between genders nor between the sanitary places (Fig. 2a).

**Figure 2.**
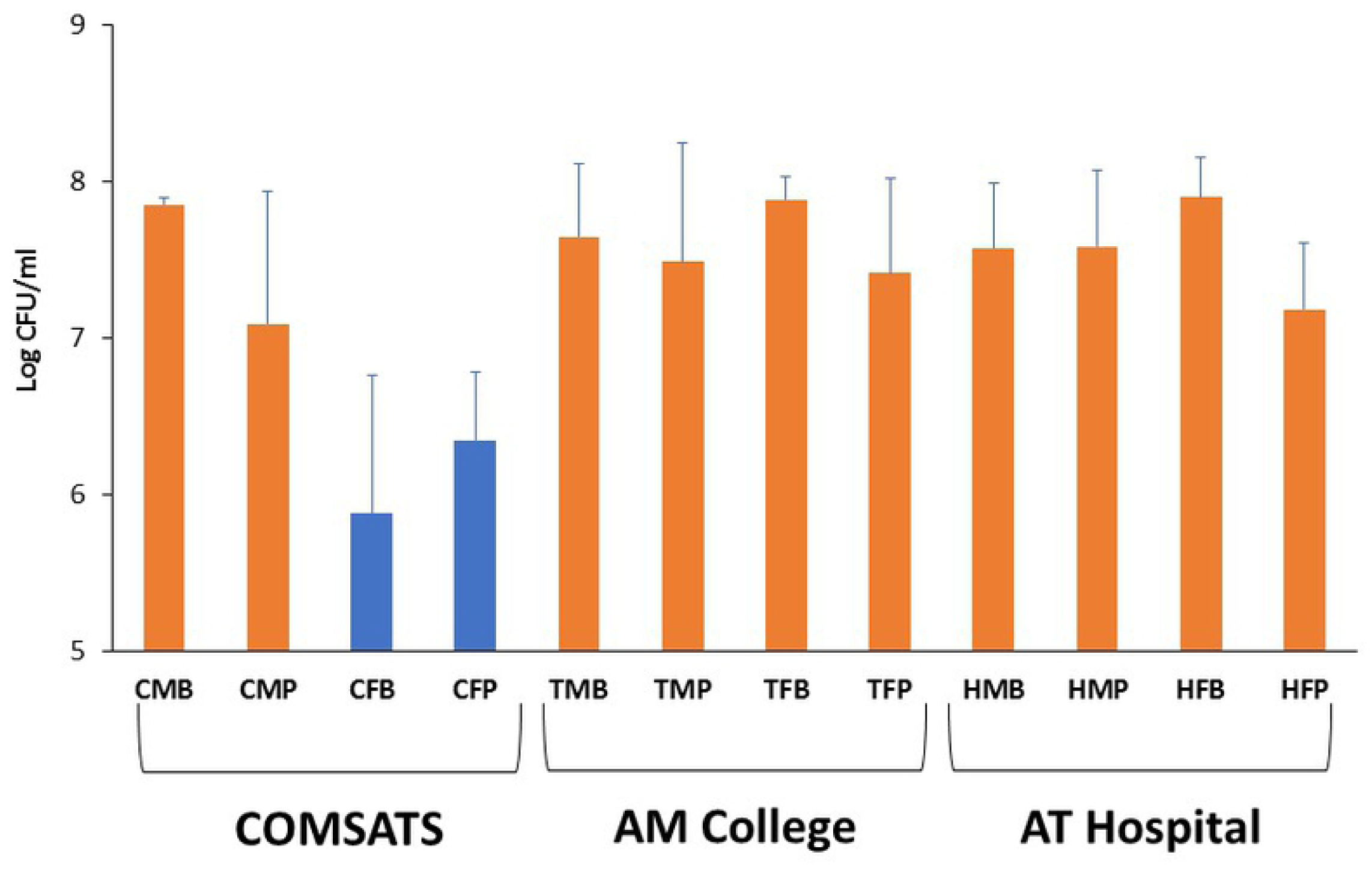
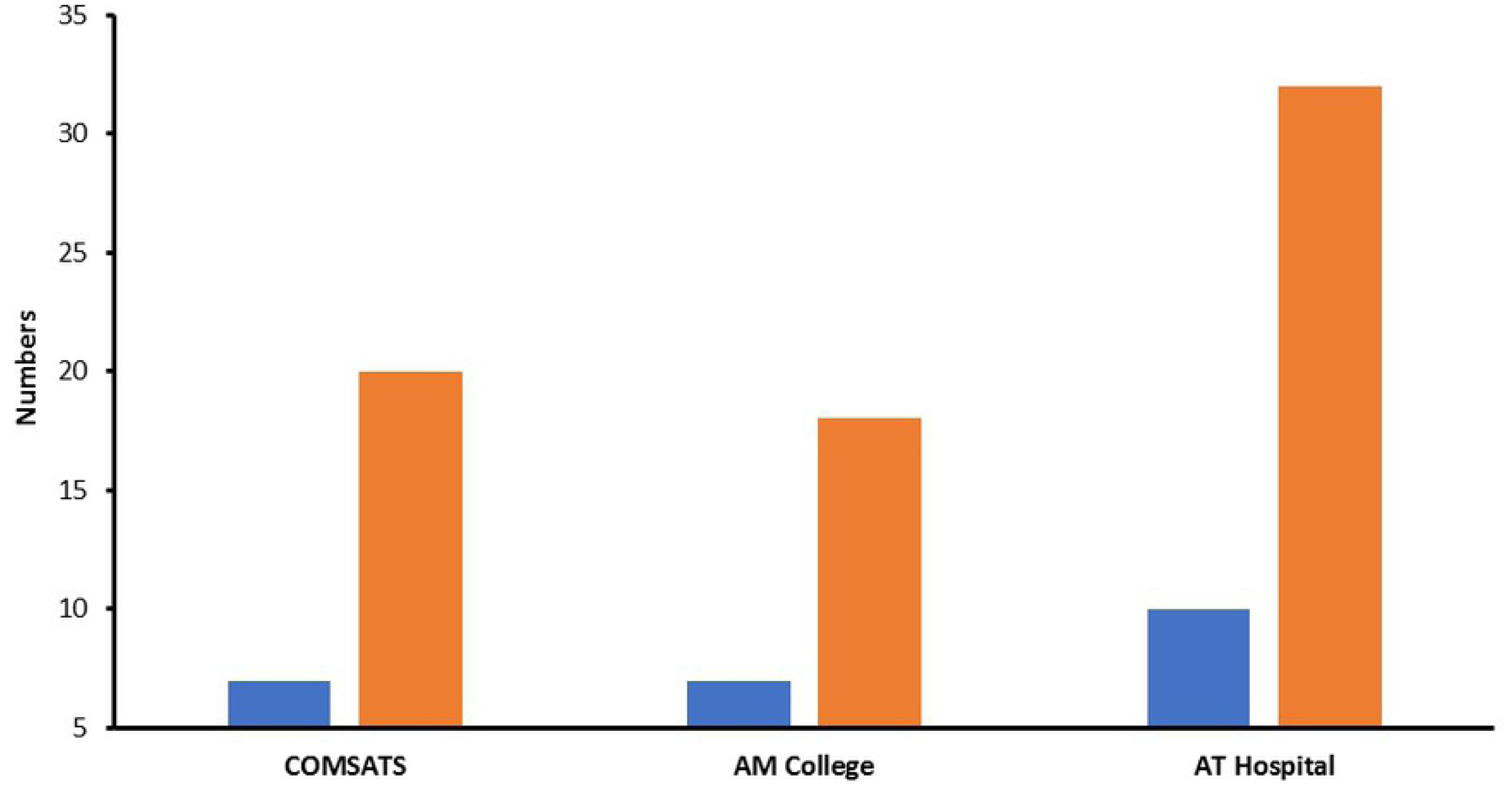
Culturable bacterial load present in different sampling points of populated human workplaces. (a) represents the colony forming units per ml for multiple places while (b) indicates bacterial morphotypes (Blue bars) observed at each workplace and the the number of bacterial strains (Orange bars) selected per site for further analyses. Error bars in (a) represent the standard deviations (n = 3) while (b) represents the absolute numbers, thus, does not have error bars.

Along with this, all the cultured bacteria were observed for varying colony morphology based on the colony shape, margins, pigmentation, and elevation from the agar surface. Based on these criteria, twenty-four different bacterial morphotypes were observed (as briefed in supplementary table 1). Out of these 24, maximally three different morphotypes were observed in a single sample, while four different samples showed this diversity of 3- morphotypes (Supplementary Figure 1). Interestingly three out of these four (i.e., CMP, HMB, HMP, and HFP) bacterially diverse samples were of the hospital, male and sanitary pot origin. On the contrary, four samples (i.e., CFB, CFP, TFB, and HFB) showed only one bacterial morphotype, and noticeably, all four were of female origin having three samples from washbasins and two from COMSATS origin (Supplementary Figure 1). For further bacterial analyses, seventy different bacterial strains were purified on the basis of varying morphotypes (i.e., approximately three bacteria from each morphotype of 24 in total). Samples with diversified cultivable bacteria were chosen for the purification of 8-12 bacterial strains from each, while 2-3 bacteria were selected from monomorphic sanitary samples (Supplementary Figure 1). Out of three sampled locations, ATH-hospital showed maximum diversity (ten different morphotypes) and so was the bacterial numbers selected from this location while other two locations were comparable for microbial diversity and selected number of bacteria for further assessments (Fig. 2b).

### Antibiotic resistance profile of the sanitation related bacterial strains

The antibiotic resistance profile of all purified 70 bacterial strains, were determined (using their resistance or sensitivity assays) via culturing the bacteria separately on the LBA medium containing various antibiotics of four different chemical classes (as detailed in Table 1). Out of seventy tested bacterial strains, 71% were resistant to Co-trimoxazole followed by Ampicillin (i.e., 67%) resistance, leaving only 29% and 33% tested bacteria to be sensitive for these antibiotics. While only 13% and 14% bacterial strains were resistant to and the rest (i.e., 87% and 86% respectively) were sensitive to Amikacin and Azithromycin respectively, which means that these are the most effective antibiotics against these tested bacterial strains (Fig. 3a) while the other four tested antibiotics exhibited the mediocre effect (approximately 60% sensitivity) for the sanitation-related bacterial strains.

**Figure 3.**
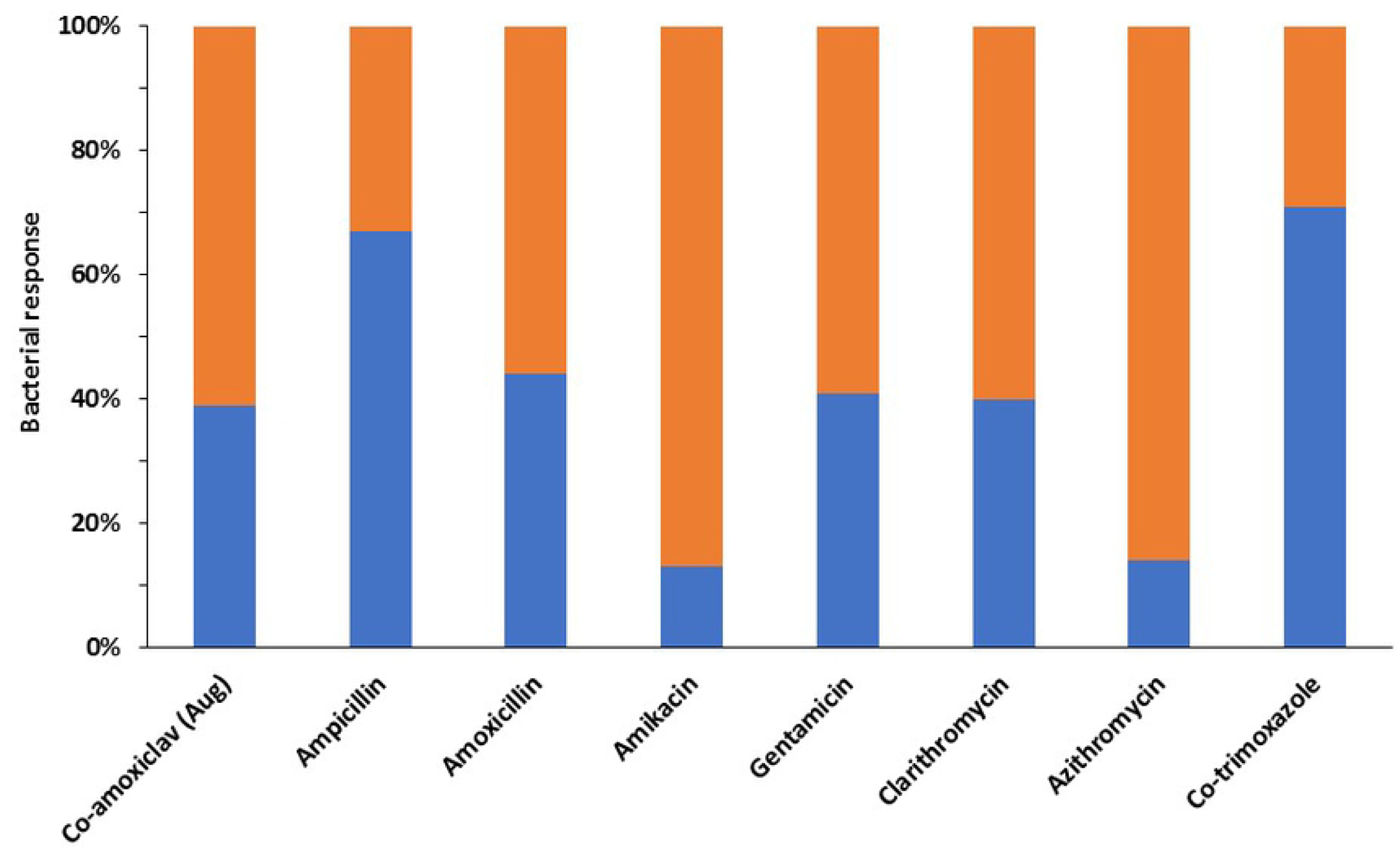
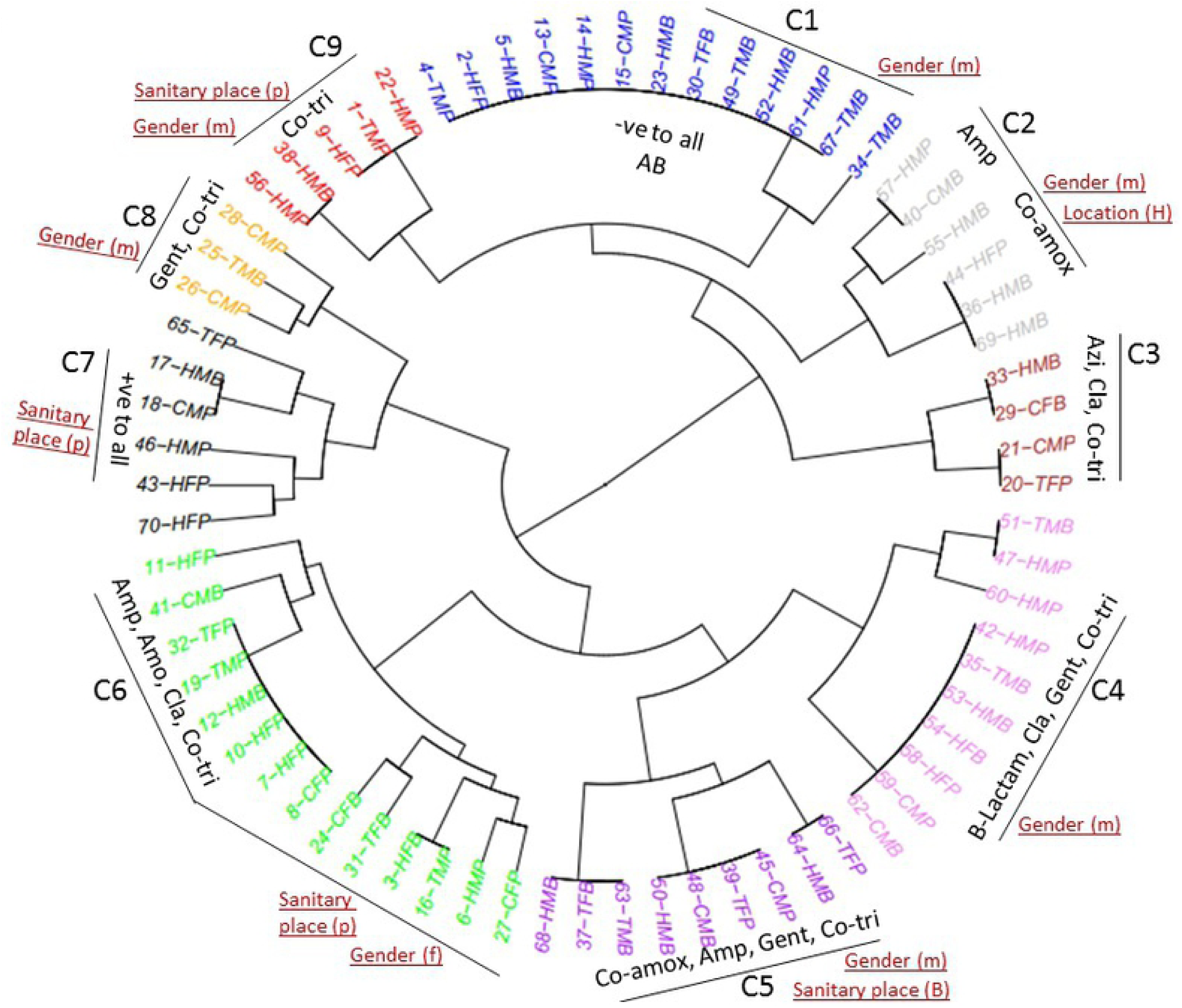
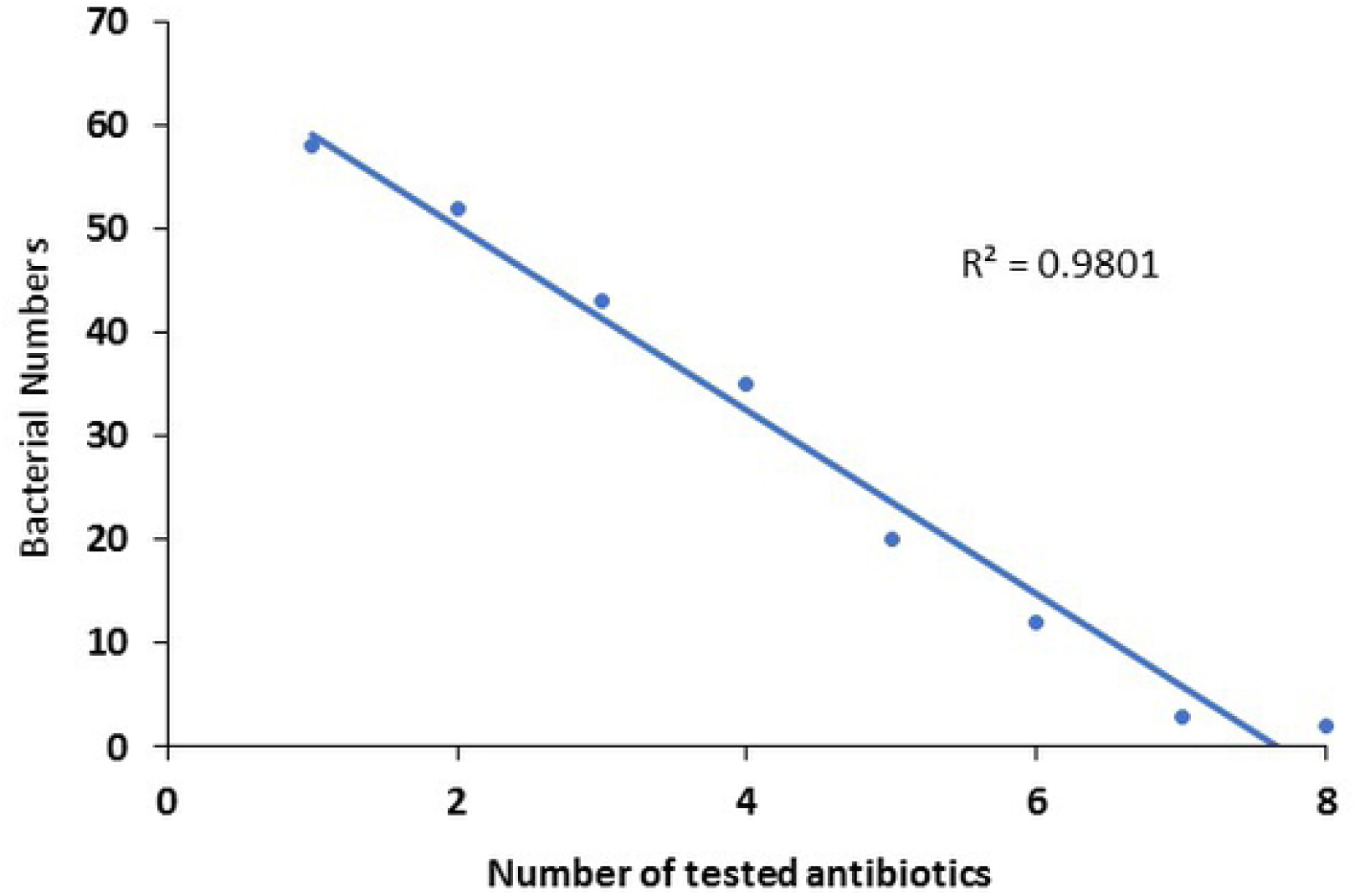
Antibiotic resistance profile of bacterial strains isolated from hygiene related environments of the populated human workplaces. (a) represents the percent resistant (Blue) and sussceptible (Orange) bacterial strains (n=70) tested for eight different antibiotics; (b) represents the antibiotic resistance profile of all tested bacterial strains grouped in various clusters based on their differential potential to resist various antibiotics; (c) indicates the multiple drug resistance (MDR) potential of purified bacterial strain against varied number of tested antibiotics. In (b) C, cluster; red colored text indicates the bacterial sampling origin potentially infuencing the AR profile clustering.

Moreover, the resistance profile of 70-bacterial strains was transformed into a numerical matrix, and “R studio” analysis software was used to analyze the grouping pattern of tested bacterial strains (Fig. 3b). The bacterial strains showing resistance to the same group of antibiotics were included in the same cluster, and so, in this way, nine different clusters were formed. The first cluster contained the bacterial strains (13 in number), which were sensitive to all the tested antibiotics. Interestingly, 11 out of 13 strains were isolated from male washroom samples, 7/13 are from washbasin, while 6/13, 5/13, and 2/13 were from the hospital, college, and COMSATS, respectively. On the contrary, the cluster-7 consists of six bacterial strains, resistant to all tested antibiotics. Noticeably, five out of six bacteria were isolated from sanitary pot samples while 4/6, 1/6, and 1/6 belonged to the hospital, college, and COMSATS respectively, while gender showed no specific role here (Fig. 3b).

The largest cluster in this analysis was the group-6 containing fourteen bacterial strains resistant to ampicillin, amoxicillin, clarithromycin, and co-trimoxazole. Out of these fourteen, nine strains were isolated from the sanitary pot and five from washbasin (similar distribution is for female and male gender respectively) while 6/14, 4/14, and 4/14 bacterial strains originated from the hospital, COMSATS and college respectively (Fig. 3b).

All other clusters showed varying antibiotic resistance potential of different tested bacterial strains. For instance, the cluster-2 showed resistance to β-lactam antibiotics. The cluster-3 consists of bacterial strains, showing resistance to azithromycin and co-trimoxazole. The fourth cluster consists of seven bacterial strains that show resistance to all the β-lactam antibiotics, clarithromycin, gentamicin, and co-trimoxazole. Cluster-5 contains 9 bacterial strains resistant to two β-lactam antibiotics, gentamicin and co-trimoxazole. The number 8 cluster showed resistance to gentamicin and co-trimoxazole. And, cluster-9 contains three bacterial strains resistant to only Sulphonamide.

If we evaluate these clusters for ARBs relevance with their isolation source (origin), majority (7/9) are derived by the gender and some (4/9) are influenced by the sampled sanitary place (mostly pot) while the sampling location seems to have mere effect on ARB grouping, based on their antibiotic resistance profiles (Fig. 3b).

### Multi-Drug Resistant Hygiene related Bacterial Strains

As mentioned above, the majority of the tested hygiene-related bacterial strains were resistant to multiple antibiotics used in this work (Supplementary table 2). Noticeably, the number of resistant bacteria decreased in number as the number of tested antibiotics was increased (Fig. 3c) that showed the inverse proportionate relationship between the both. Based on these observations, 23-bacteria were selected to further evaluate for identification and plasmid curing purposes. Out of these 23 multi-drug resistance (MDR) bacterial strains, nine strains were from Ayub Teaching Hospital, seven strains each from Ayub Medical College, and COMSATS University (Table 2). Of the nine multi-drug resistant bacterial strains isolated from ATH, seven strains were from male washroom samples, and five strains were from washroom basin samples. Out of the seven MDR bacterial strains isolated from AMC samples, four belongs to female washroom samples and washroom sanitary pot samples, while the seven MDR bacterial strains isolated from COMSATS University contain five strains from the male washroom and washroom sanitary pot samples.

**Table 2:** Characteristics of purified bacterial strains and their 16s rRNA based molecular identification

| Serial No. | Strain code | Origin |  |  |  |  |  |  | No. of Antibiotics |  | Nearest hit (NH) | Identity | Accession number of NH |
| --- | --- | --- | --- | --- | --- | --- | --- | --- | --- | --- | --- | --- | --- |
|  |  | Site |  |  | Gender |  | Place |  |  |  |  |  |  |
|  |  | C | T | H | M | F | B | P | R | S |  |  |  |
| 1 | HFP.7 |  |  |  |  |  |  |  | 4 | 4 | *Pseudomonas putida* MHT-PSE-01 | 98% | MH279662.1 |
| 2 | TMB.11 |  |  |  |  |  |  |  | 5 | 3 | *Pseudomonas oryzihabitans* ML-25-4 | 98% | KJ401064.1 |
| 3 | CMP.16 |  |  |  |  |  |  |  | 3 | 5 | *Pseudomonas putida* AGL 13 | 100% | EU118779.1 |
| 4 | HMP.17 |  |  |  |  |  |  |  | 8 | 0 | *Pseudomonas oryzihabitans* FP42 | 99% | MH620723.1 |
| 5 | CMP.19 |  |  |  |  |  |  |  | 4 | 4 | *Stenotrophomonas rhizophila* | 99% | FM178869.1 |
| 6 | HMB.21 |  |  |  |  |  |  |  | 3 | 5 | *Acinetobacter lwoffii* 2A | 98% | KF993657.1 |
| 7 | TFP.24 |  |  |  |  |  |  |  | 5 | 3 | *Pseudomonas japonica* 58F5 | 99% | MG269716.1 |
| 8 | CMP.25 |  |  |  |  |  |  |  | 4 | 4 | *Rheinheimera* sp. 193 | 100% | JQ012969.1 |
| 9 | HMP.26 |  |  |  |  |  |  |  | 5 | 3 | *Comamonas denitrificans* 14 | 99% | DQ836252.1 |
| 10 | CFB.28 |  |  |  |  |  |  |  | 4 | 4 | *Delftia* sp. D6 | 99% | MG594845.1 |
| 11 | CFP.31 |  |  |  |  |  |  |  | 4 | 4 | *Bacillus cereus* LAHAAB_29 | 99% | KX908030.1 |
| 12 | TFB.35 |  |  |  |  |  |  |  | 6 | 2 | *Pseudomonas* sp. WS14 | 97% | MG807361.1 |
| 13 | TFP.40 |  |  |  |  |  |  |  | 1 | 7 | *Acinetobacter johnsonii* | 98% | MH636837.1 |
| 14 | TMP.46 |  |  |  |  |  |  |  | 6 | 2 | *Sphingobacterium alimentarium* | 99% | FN908504.1 |
| 15 | HMB.47 |  |  |  |  |  |  |  | 5 | 3 | *Pseudomonas putida* strain HTc1 | 99% | JF703647.1 |
| 16 | CMB.51 |  |  |  |  |  |  |  | 5 | 3 | *Pseudomonas fluorescens* FC6846 | 98% | MH497588.1 |
| 17 | CMP.55 |  |  |  |  |  |  |  | 2 | 6 | *Bacillus thuringiensis* DST212 | 99% | MH793408.1 |
| 18 | HMP.60 |  |  |  |  |  |  |  | 4 | 4 | *Pseudomonas putida* PP | 98% | MH368654.1 |
| 19 | HMB.64 |  |  |  |  |  |  |  | 5 | 3 | *Bacillus anthracis* USW-ERY-2 | 99% | MF083049.1 |
| 20 | TFP.65 |  |  |  |  |  |  |  | 7 | 1 | *Aeromonas caviae* Moschidae | 99% | KX648715.1 |
| 21 | TMB.67 |  |  |  |  |  |  |  | 4 | 4 | *Pseudomonas putida* D19 | 99% | EF204247.1 |
| 22 | HMB.69 |  |  |  |  |  |  |  | 2 | 6 | *Brevundimonas diminuta* 264AG7 | 97% | KF836539.1 |
| 23 | HFP.70 |  |  |  |  |  |  |  | 6 | 2 | *Brevundimonas terrae* STM25 | 99% | KY393019.1 |
|  |  | 7 | 7 | 9 | 15 | 8 | 9 | 14 |  |  |  |  |  |

\*Green color indicates the origin of bacteria. C, COMSATS; T, medical teaching college; H, hospital; M, male; F, female; B, washbasin; P, sanitary pot.

In regards to the molecular analysis of MDR, the earlier work has reported the presence of tetracycline resistance genes in bacterial species isolated from wastewater samples [25]. Therefore, the qualitative presence of tetracycline-resistant genes i.e., *Tet* A and *Tet* M was also evaluated (PCR via gene-specific primers) in our sanitation-related MDR bacterial strains. And, only one strain i.e., *Pseudomonas putida* CMP-16, isolated from male washroom samples of COMSATS, was found positive for these genes.

### Molecular and Phylogenetic analyses of sanitation related MDR bacteria

Subsequent to DNA extraction from 23 MDR bacterial strains, a 16s rRNA gene-based PCR was performed via using the universal bacterial primers (i.e., 27F and 1492R). After the amplification of the 16s rRNA gene, samples were run on 1% gel electrophoresis, and amplification was confirmed by specified band size. Afterwards, the PCR products were sent for DNA sequencing.

On the receipt, the sequence data were manually curated with Chromas and BLAST analyzed on the online NCBI database. The reference sequences, similar to the hygiene-related 23 bacterial sequences, along with percent identity, are listed in table 2. The majority of the sequences of sanitation-related bacterial isolates shared 99-100% homology with already reported bacterial sequences (Table 2).

Moreover, a phylogenetic tree was constructed to exhibit the evolutionary relationship of different bacteria with each other. For this purpose, subsequent to the sequence alignment of 23 hygiene-related bacterial strains and of reference sequences, a neighbor-joining phylogenetic analysis was performed, and resultant tree grouped our 23 bacterial strains into various taxonomic clusters (Fig. 4). For instance, the bacterial strains HMP-17, TMB-67, CMP-16, and TMB-11 clustered with *Pseudomonas oryzihabitans* and *P. putida,* strains that were isolated from soil and industrial wastewater having similar origin like the antibiotic resistance strains we isolated. Interestingly, all these four bacterial strains were isolated from male washroom samples. Another group of *P. putida* strains includes HFP-7, TFB-35, and HMB-47, which were originated from ATH/AMC while the reference sequences are from wastewater, lake sediment samples, and river water. TFP-24 and HMP-60, both strains isolated from sanitary pot samples, are clustered with *P. japonica* and *P. putida* strains, also isolated from wastewater and wastewater treatment plant. CMB-51 is grouped with *P. fluorescens* and *P. gessardii* strains isolated from patients, WWTP, and sludge.

**Figure 4.**
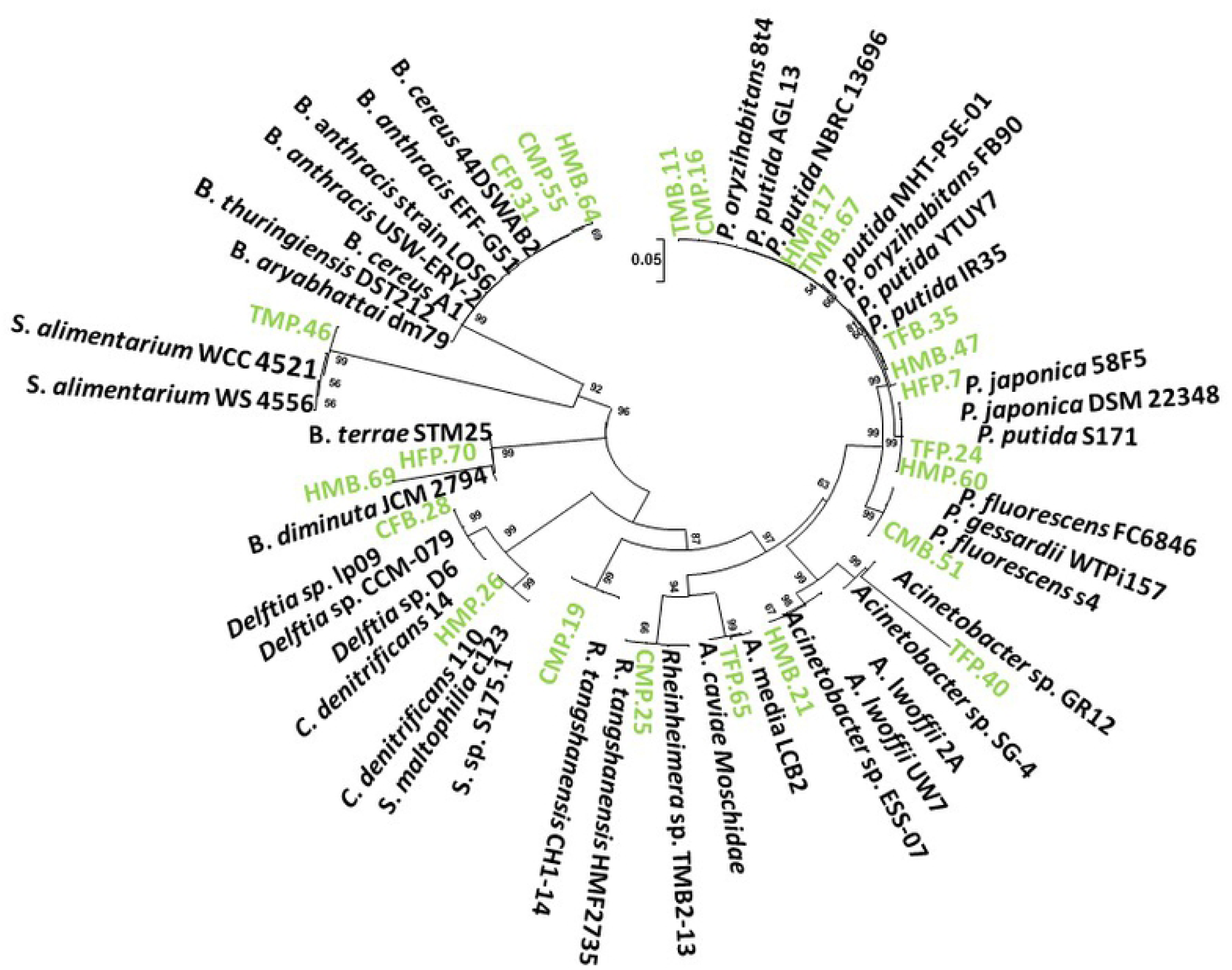
Phylogenetic analysis of the purified antibiotic resistant bacterial strains, based on their 16s rRNA gene sequences. The tree is constructed via neighbor joining method considering Maximum Composite Likelihood model with Bootstrap replicates of 1000, and bootstrap values equal to or greater than 50 are shown.

TFP-40 strain is grouped with *Acinetobacter sp.* having their sources from lake sediments and plant roots. Whilst, HMB-21 and TFP-65 are grouped with *A. lwoffii* and *Aeromonas* strains, which have similar isolation sources like of our strains i.e., human skin, wastewater, and WWTP. CMP-19 is clustered with *Rheinheimera sp*., while HMP-26 and CFB-28 have evolutionary closeness with *Comamonas denitrificans* and *Delftia sp.* strains isolated from sludge waste, WWTP, wastewater sludge and contaminated water. HMB-69 and HFP-70 have an evolutionary relationship with *Brevundimonas diminuta* and *B. terrae* strains. TMP-46 shared a common cluster with *Sphingobacterium alimentarium* strains. Noticeably, the hygiene-related bacterial strains CFP-31, CMP-55, and HMB-64 have been clustered with Gram-positive *Bacillus* strains (Fig. 4).

### Plasmid Curing and Antibiotics Resistance Profiling of hygiene related MDR Bacteria

Plasmid curing experiment was performed for (34) bacterial strains that were resistant to multiple antibiotics. After 5-days of bacterial culture incubation (along with plasmid replication inhibitor i.e. acridine orange), 10 out of 34 bacterial strains (i.e., HFB-3, HMP-6, CMP-26, HMB-36, TFB-37, CMB-40, HMP-42, HFP-44, CMP-45 and HFB-54) lost their plasmids. More specifically, 5.9% of the bacteria were found cured after 2-days and further 23.5% of the strains got cured after 5-days of incubation. While the further incubation did not produce anymore curing.

After the plasmid curing, these MDR bacterial strains were re-evaluated for their resistance potential against tested antibiotics (Fig. 5). The bacterial strain HFB-3 was ampicillin-resistant before plasmid curing, but it lost this resistance after plasmid curing. CMP-26 strain was resistant to amikacin and gentamicin, but plasmid curing made it sensitive for both. Similarly, the TFB-37 strain was resistant to ampicillin and gentamicin, which were lost due to plasmid curing. Strain HMP-42 was also resistant to co-amoxiclav and gentamicin but not anymore after plasmid curing. HFP-44 resistance to ampicillin was also diminished by its plasmid loss. CMP-45 was also no longer resistant to gentamicin. Similarly, the resistance of HFB-54 to co-amoxiclav and amoxicillin was also lost after plasmid curing (Fig. 5).

**Figure 5.**
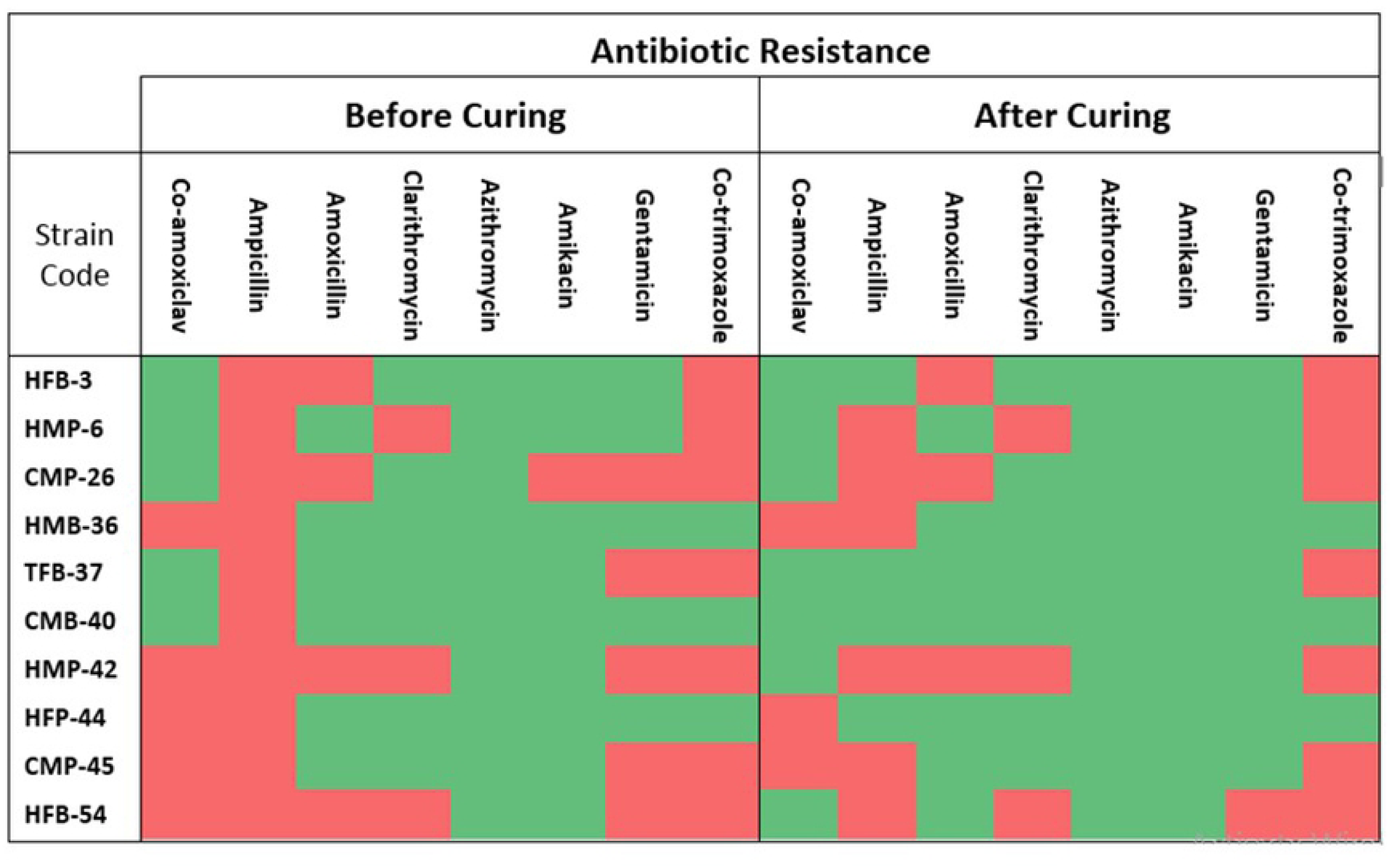
Pre and post plasmid-curing antibiotic resistance profile of isolated bacterial strains, depicting their antibiotic resistance conduit potentially on their respective plasmids. The post plasmid-curing loss of antibiotic resistance potential indicates the likely presence of antibiotic resistant gene(s) on the cured (lost) plasmid for that particular bacteria.

The evidence that bacterial strains lost their resistance to tested antibiotics after plasmid curing may corroborate that the antibiotic resistance genes in these bacterial strains were present on their respective plasmids. While the bacterial strains, resistant to the antibiotics before and even after plasmid curing experiment, may have their resistance genes present on their genomic DNA or on other incompatible plasmids.

## Discussion

This study has assessed the prevalence of multi-drug resistant bacteria in various hygiene related conditions prevailing in populated human workplaces. The unhygienic environment normally serves as a reservoir for antibiotic resistance in different bacteria through HGT, leading to environmental multi-drug resistance. In these reservoirs, different pathogenic microbes, if present, may cause different infectious diseases in humans [26]. In this regard, our results suggest that the bacterial strains isolated from AMC-ATH show resistance to similar antibiotics, meaning that they may have transferred their AR genes to each other. Both these sample-locations are present in close proximity to each other, and human flow from one location to another is very common meaning that the transfer of bacteria from one location to another might be through human movement, which may ultimately pose the concerns for ARGs transfer to other bacterial hosts.

Bacterial numbers were observed quite high in our sanitation-related samples i.e., up to 10^8^ CFU/ml in majority samples. Moreover, the bacterial morphological and taxonomical diversity was also very high, which means both of these factors (abundance and diversity) may increase the chances of resistance genes transfer through HGT between different bacterial species. This has earlier been reported that ARGs are abundantly present in wastewater and increase the chances of antibiotic resistance in bacteria present in there [27]. The bacteria may also adopt and mutate their genome due to the selection pressure of antibiotics present in their environment.

Antibiotic resistance profiling was performed for hygiene-related bacterial strains, isolated from various places of washrooms. The samples from male and female washrooms were separately evaluated, and the results showed that female washroom samples from COMSATS have significantly fewer bacteria as compared to the male washrooms of the same location. This is consistent with previous literature where statistically significant differences were observed between male and female samples for *E. coli* susceptibilities against different antibiotics [28]. Recently, a 10-years study of Portuguese patients’ also reported the difference in antibiotic resistance by patient sex [43]. The researchers reported that urinary *E. coli* isolates from the male were more resistant to their tested antibiotics as compared to female-origin isolates [43]. Yet, another study reported their male samples at a higher risk for selecting antimicrobial resistance [29].

Beyond the gender influences on bacterial community establishment and AR development, the populated human workplaces may even offer the possibilities of AR dissemination; these places being the hotspots for human gatherings, interactions and subsequently may serve as the source for microbial pileup and genetic shift over between different bacteria. Our results have shown that the workplaces with enhanced human flow have increased risk of ARB presence, and therefore the transfer of ARGs may significantly be more in such places. According to a report, antibiotics used in agriculture, can still be found in dairy, pig, poultry and factory farms and may travel into the municipal wastewater and even contaminate the groundwater, hence promoting antibiotic resistance [44]. Humans may also get infected with ARB via exposure to contaminated environments. Different studies have earlier shown that European broiler farmers and turkey farmers, as well as other workers, were at increased risk of colonization with antimicrobial resistant *E. coli* and *Enterococcus* because of their occupational exposure to these contaminated environments [30]. A study performed in the Netherlands showed that MRSA infections were estimated to be 29% in pig farmers as compared to the general population where it was only 0.1% [31]. Similarly, hospitals and intensive care units also contribute greatly to the development and spread of ARB [32]. The health care staff and attendants are at greater risk of ARB because they are in contact with patients infected with different bacteria, and with surfaces like door handles, contaminated equipment’s and patients’ body fluids [33].

Bacterial strains identified in this study showed resistance to multiple antibiotics. It has been reported in many earlier studies that *Pseudomonas putida* is pathogenic under certain conditions, can infect humans and may become resistant to many antibiotics [34]. Similarly, our data that exhibited the resistance potential of *Pseudomonas putida* to β-lactam, macrolides, and Sulphonamide antibiotics. *Acinetobacter lwoffii* strains cause nosocomial infections in humans [9], and our *Acinetobacter* strains are resistant to multiple antibiotics that demonstrate the potential relevance with human-inhabited environments. Similarly, *Bacillus cereus* is well-known pathogen causing diarrhea, food poisoning to humans and harbors a variety of plasmids carrying ARGs with transfer potential to other bacterial species, thus, playing a role in the prevalence of MDR environmental bacterial strains [35]. In this regard, our *Bacillus* strains showed resistance to macrolide antibiotics, and *Bacillus anthracis* has already been reported to causes anthrax infection [36]. Interestingly, our three *Bacillus* strains (HMB64, CMP55, and CFP31) have different sources of isolation, though COMSATS, male washroom, and the sanitary pot is common in two of these three strains. These three bacteria were resistant to ampicillin and sensitive to azithromycin and amikacin with differentiating profiles for the other five tested antibiotics. These characteristics, along with taxonomic identity, indicate their common AR strategy, which probably has been evolved at some interception (e.g., sanitation) where their isolation sources may mix up.

Moreover, *Brevundimonas diminuta* has become a major opportunistic pathogen associated with nosocomial and urinary tract infections. These species have the ability to pass even through sterilizing filters, which may reinforce their potential for harmful infections [37]. Noticeably, our hygiene-related bacterial collection contains two MDR tentative *Brevundimonas* isolates (HMB69 and HFP70) of hospital origin and, therefore, the disease prevention programs should consider temporal investigations for *Brevundimonas* spp. monitoring and possible outbreaks as these bacteria are of severe clinical significance [38]. *Stenotrophomonas maltophilia,* on the other hand, is highly pathogenic and can cause serious diseases, sometimes non-treatable, in individuals with weak immunity [39], with its intrinsic resistance to a number of antibiotics [40]. *S. maltophilia* has been reported as resistant to different sulphonamide antibiotics that is in line with our results showing *S. maltophilia* CMP19 as resistant to sulphonamide and other antibiotics tested here.

Earlier studies have also demonstrated that the antibiotic resistance in bacteria is usually present on plasmids, which transfer to other bacteria through HGT, and thus these plasmids are the real players for AR spread [41]. In our study, plasmid curing abolished the resistance of different cured bacterial strains against some antibiotics, which suggests that ARGs in our tested bacteria are also present on plasmids. For instance, *Comamonas denitrificans* strain HMP-26 and *Acinetobacter johnsonii* strain TMP-40 were originally resistant to amikacin, gentamicin, and ampicillin, but this resistance was diminished after plasmid curing. A similar study performed by [42] demonstrated that plasmid curing decreased the bacterial resistance to β-lactams, aminoglycosides, and tetracycline antibiotics [42].

Our results also showed that bacteria having similar AR patterns cluster together for their isolation source as well (Figure 3b), e.g., male/female washroom samples, sanitary pot/basin, and sampling workplaces. Some bacterial strains isolated from ATH (hospital) showed resistance to similar antibiotics such as strains HFP-44, HMP-36, HMB-69 are resistant to azithromycin, and Sulphonamide. This may indicate that bacterial strains showing resistance to similar antibiotics, and also isolated from the same location, may have acquired resistance from the same source, probably, through HGT. Similarly, bacterial strains TMB-51 and HMP-47 are both resistant to β-lactam antibiotics, sulphonamides and clarithromycin, while both strains are originated from the male washroom source, and their presence in medical college and teaching hospital indicates the possibility of bacterial transfer from hospital to college or vice versa which would have happened through human (students) movement in their workplaces.

From our data, it has also been observed that amikacin (aminoglycoside) is the most effective antibiotic as only 9 bacteria strains (out of 70) were resistant while Co-trimoxazole (Sulphonamide) was the least effective with 50 resistant strains. Therefore, Co-trimoxazole resistance seems rapidly developing that actually is an alarming situation for human health. Interestingly, the bacterial abundance and diversity in female washroom samples (particularly in COMSATS) were considerably less as compared to the respective male washroom samples. The possible reason for this observation could be the hygienic sense of females or the likely use of cosmetic items, which may inhibit bacterial growth (e.g., via heavy metal supplementation in makeup products). It has also been observed that 22 bacterial strains isolated from female washroom samples were mostly resistant to β-Lactam antibiotics. Moreover, we plan in future to extend the sampling size to several other workplaces with diversified sampling locations like offices etc. along with extended spectrum of test antibiotics. On parallel, the molecular analyses e.g. quantification of different ARGs and mobile genetic elements are also expected to be included in future research.

## Conclusion

It may be concluded that the hygiene-related sanitary conditions contribute greatly in occurrence and potential spread of antibiotic-resistant bacteria. Bacterial strains isolated from washroom samples of human workplaces showed resistance to multiple antibiotics that are currently in clinical use. Such increased number of multiple antibiotic resistant bacteria will have a very negative impact on human health in future. Therefore, it is the need of time to overcome this bacterial resistance. And, the limited use of antibiotics is one option so that the bacteria are not put on pressure to adopt or acquire the antibiotic resistance. And, there is also a need to improve our hygienic conditions and sanitation facilities to decrease the bacterial growth in these environments and consequently to alleviate the chances of infection prevalence within the human workplaces.

## Acknowledgement

This scientific compilation was financially supported by research grants from Higher Education Commission (HEC) of Pakistan (NRPU20-3657/R&D/HEC/14/704) and TWAS-COMSTECH (18-363 RG/REN/AS_C – FR3240305796). Kinza Irshad is acknowledged to assist in sampling, particularly in female facilities.

## References

1. Bouki C, Venieri D, Diamadopoulos E. Detection and fate of antibiotic resistant bacteria in wastewater treatment plants: A review. Ecotoxicol Environ Saf. 2013;91: 1–9. doi: 10.1016/j.ecoenv.2013.01.016

2. Béahdy J. Recent developments of antibiotic research and classification of antibiotics according to chemical structure. Adv Appl Microbiol. 1974;18: 309–406. doi: 10.1016/S0065-2164(08)70573-2

3. Wright GD. Antibiotic resistance : where does it come from and what can we do about it? BMC Biology. 2010; 8(1): 123.

4. Van Hoek AHAM, Mevius D, Guerra B, Mullany P, Roberts AP, Aarts HJM. Acquired antibiotic resistance genes: An overview. Front Microbiol. 2011;2: 1–27. doi: 10.3389/fmicb.2011.00203

5. Davies J, Davies D. Origins and evolution of antibiotic resistance. Microbiol Mol Biol Rev. 2010;74: 417–433. doi: 10.1128/MMBR.00016-10

6. Sommer MOA, Munck C, Toft-Kehler RV, Andersson DI. Prediction of antibiotic resistance: Time for a new preclinical paradigm? Nat Rev Microbiol. 2017;15: 689–696. doi: 10.1038/nrmicro.2017.75

7. Andersson DI. Persistence of antibiotic resistant bacteria. Curr Opin Microbiol. 2003;6: 452–456. doi: 10.1016/j.mib.2003.09.001

8. Schwartz T, Kohnen W, Jansen B, Obst U. Detection of antibiotic-resistant bacteria and their resistance genes in wastewater, surface water, and drinking water biofilms. FEMS microbiology ecology. 2003;43(3): 325–35.

9. Hu R, Liao S, Huang C, Huang Y, Yang T. An inducible fusaric acid tripartite efflux pump contributes to the fusaric acid resistance in Stenotrophomonas maltophilia. PLoS One 2012;7: 1–8. doi: 10.1371/journal.pone.0051053

10. Versluis D, D’Andrea MM, Ramiro Garcia J, Leimena MM, Hugenholtz F, Zhang J, et al. Mining microbial metatranscriptomes for expression of antibiotic resistance genes under natural conditions. Sci Rep. 2015;5: 1–10. doi: 10.1038/srep11981

11. Blair JMA, Webber MA, Baylay AJ, Ogbolu DO, Piddock LJ V. Molecular mechanisms of antibiotic resistance. Nat Rev Microbiol. 2015;13: 42–51. doi: 10.1038/nrmicro3380

12. Spellberg B, Guidos R, Gilbert D, Bradley J, Boucher HW, Scheld WM, et al. The epidemic of antibiotic-resistant infections: A call to action for the medical community from the Infectious Diseases Society of America. Clin Infect Dis. 2008;46: 155–164. doi: 10.1086/524891

13. Walsh C. Molecular mechanisms that confer antibacterial drug resistance. Nature. 2000;406: 775–781.

14. Yang Q, Tian T, Niu T, Wang P. Molecular characterization of antibiotic resistance in cultivable multidrug-resistant bacteria from livestock manure. Environ Pollut. 2017;229: 188–198. doi: 10.1016/j.envpol.2017.05.073

15. San Millan A, Escudero JA, Gifford DR, Mazel D, MacLean RC. Multicopy plasmids potentiate the evolution of antibiotic resistance in bacteria. Nat Ecol Evol. 2016;1: 0010. doi: 10.1038/s41559-016-0010

16. Allen HK, Donato J, Wang HH, Cloud-Hansen KA, Davies J, Handelsman J. Call of the wild: Antibiotic resistance genes in natural environments. Nat Rev Microbiol. 2010;8: 251–259. doi: 10.1038/nrmicro2312

17. Pitout JDD. Extraintestinal pathogenic Escherichia coli: A combination of virulence with antibiotic resistance. Front Microbiol. 2012;3: 1–7. doi: 10.3389/fmicb.2012.00009

18. Mellata M. Human and avian extraintestinal pathogenic Escherichia coli : infections, zoonotic risks, and antibiotic resistance trends. Foodborne Pathogens and Disease. 2013;10(11): 916–32. doi: 10.1089/fpd.2013.1533

19. Nazir R, Warmink JA, Voordes DC, Bovenkamp HH Van De, Elsas JD Van. Inhibition of mushroom formation and induction of glycerol release — Ecological strategies of Burkholderia terrae BS001 to create a hospitable niche at the fungus Lyophyllum sp . Strain Karsten. Microb. Ecol. 2013;65(1): 245–54. doi: 10.1007/s00248-012-0100-4

20. Mukherjee A, Ghosh K. Antagonism against fish pathogens by cellular components and verification of probiotic properties in autochthonous bacteria isolated from the gut of an Indian major carp, Catla catla (Hamilton). Aquacult. Res. 2014; 1–13. doi: 10.1111/are.12676

21. Nazir R, Shen JP, Wang JT, Hu HW, He JZ. Fungal networks serve as novel ecological routes for enrichment and dissemination of antibiotic resistance genes as exhibited by microcosm experiments. Sci Rep. 2017;7: 1–13. doi: 10.1038/s41598-017-15660-7

22. Lu XM, Li WF, Li C Ben. Characterization and quantification of antibiotic resistance genes in manure of piglets and adult pigs fed on different diets. Environ Pollut. 2017;229: 102–110. doi: 10.1016/j.envpol.2017.05.080

23. Janda JM, Abbott SL. 16S rRNA gene sequencing for bacterial identification in the diagnostic laboratory : Pluses, perils, and pitfalls. J. Clin. Microbiol. 2007;45: 2761–2764. doi: 10.1128/JCM.01228-07

24. Ghosh S, Mahapatra NR. Plasmid curing from an acidophilic bacterium of the genus Acidocella. FEMS Microbiol. Lett. 2000;183: 271–274.

25. Szczepanowski R, Linke B, Krahn I, Gartemann K, Gu T, Eichler W, et al. Detection of 140 clinically relevant antibiotic-resistance genes in the plasmid metagenome of wastewater treatment plant bacteria showing reduced susceptibility to selected antibiotics. Microbiology. 2009; 2306–2319. doi: 10.1099/mic.0.028233-0

26. Wellington EMH, Boxall ABA, Cross P, Feil EJ, Gaze WH, Hawkey PM, et al. The role of the natural environment in the emergence of antibiotic resistance in Gram-negative bacteria. Lancet Infect Dis. 2013;13: 155–165. doi: 10.1016/S1473-3099(12)70317-1

27. Baquero F, Martínez J-L, Cantó N R, Pruzzo C, Canepari P. Antibiotics and antibiotic resistance in water environments. Curr. Opin. Biotechnol. 2008;19: 260–265. doi: 10.1016/j.copbio.2008.05.006

28. Mcgregor JC, Elman MR, Bearden DT, Smith DH. Sex- and age-specific trends in antibiotic resistance patterns of Escherichia coli urinary isolates from outpatients. BMC Family Practice. 2013;14(1):25.

29. Cole MJ, Spiteri G, Jacobsson S, Woodford N, Tripodo F, Amato-gauci AJ, et al. Overall low extended-spectrum cephalosporin resistance but high azithromycin resistance in Neisseria gonorrhoeae in 24 European Countries. BMC Infect. Dis. 2017;17(1): 617. doi: 10.1186/s12879-017-2707-z

30. Price LB, Graham JP, Lackey LG, Roess A, Vailes R, Silbergeld E. Elevated risk of carrying gentamicin-resistant Escherichia coli among U.S poultry workers. Environ. Health Perspect. 2007; 1738–1742. doi: 10.1289/ehp.10191

31. Dutkiewicz J, Cisak E, Sroka J, Wójcik-fatla A, Zając V. Biological agents as occupational hazards – selected issues. Annals of Agricultural and Environmental Medicine. 2011;18: 286–293.

32. Brantley SJ, Argikar AA, Lin YS, Nagar S, Paine MF. Herb–drug interactions: challenges and opportunities for improved predictions. Drug Metabolism and Disposition. 2014;42(3): 301–17.

33. Struelens MJ. The epidemiology of antimicrobial resistance in hospital acquired infections : problems and possible solutions. BMJ. 1998;317: 2–4.

34. Yoshino Y, Kitazawa T. Pseudomonas putida bacteremia in adult patients : five case reports and a review of the literature. J. Infect. Dis. Chemother. 2011; 278–282. doi: 10.1007/s10156-010-0114-0

35. Chon J, Kim J, Lee S, Hyeon J, Seo K. Toxin pro file, antibiotic resistance, and phenotypic and molecular characterization of Bacillus cereus in Sunsik. Food Microbiol. 2012;32: 217–222. doi: 10.1016/j.fm.2012.06.003

36. Brook I, Elliott TB, Ii HIP, Sautter TE, Gnade BT, Thakar JH, et al. In vitro resistance of Bacillus anthracis Sterne to doxycycline, macrolides and quinolones. Int. J. Antimicrob. Agents. 2001;18: 559–562.

37. Han XY, Andrade RA. Brevundimonas diminuta infections and its resistance to fluoroquinolones. J. Antimicrob. Chemother. 2005;55(6): 853–9. doi: 10.1093/jac/dki139

38. Ryan MP, Pembroke JT, Ryan MP, Pembroke JT, Ryan MP, Pembroke JT. Brevundimonas spp : Emerging global opportunistic pathogens Brevundimonas spp : Emerging global opportunistic pathogens. Virulence. 2018;9: 480–493. doi: 10.1080/21505594.2017.1419116

39. Chang Y, Lin C, Chen Y, Hsueh P. Update on infections caused by Stenotrophomonas maltophilia with particular attention to resistance mechanisms and therapeutic options. Front. Microbiol. 2015;6: 1–20. doi: 10.3389/fmicb.2015.00893

40. Sánchez MB. Antibiotic resistance in the opportunistic pathogen Stenotrophomonas maltophilia. Front. Microbiol. 2015;6: 1–7. doi: 10.3389/fmicb.2015.00658

41. Lopatkin AJ, Meredith HR, Srimani JK, Pfeiffer C, Durrett R, You L. antibiotic resistance. Nat Commun. 2017;8(1): 1–0. doi: 10.1038/s41467-017-01532-1

42. Rosander A, Connolly E, Roos S. Removal of antibiotic resistance gene-carrying plasmids from Lactobacillus reuteri ATCC 55730 and characterization of the resulting daughter strain, L. reuteri DSM 17938. Appl Environ Microbiol. 2008;74: 6032–6040. doi: 10.1128/AEM.00991-08

43. Linhares I, Raposo T, Rodrigues A, Almeida A. Frequency and antimicrobial resistance patterns of bacteria implicated in community urinary tract infections: a ten-year surveillance study (2000–2009). BMC infectious diseases. 2013;13(1): 19. doi.org/10.1186/1471-2334-13-19

44. World Health Organization. Antimicrobial resistance: global report on surveillance. World Health Organization; 2014.

45. Tennstedt T, Szczepanowski R, Braun S, Pühler A, Schlüter A. Occurrence of integron-associated resistance gene cassettes located on antibiotic resistance plasmids isolated from a wastewater treatment plant. FEMS Microbiology Ecology. 2003;45(3): 239–52. doi.org/10.1016/S0168-6496(03)00164-8

